# Role of nuclear ATPases in nuclear mechanics and cell migration through confined spaces: opposite effects of BRG1 and cohesin

**DOI:** 10.1101/2025.09.21.677586

**Authors:** Łukasz Suprewicz, Fitzroy J. Byfield, Thomas T. Dutta, Paul A. Janmey

## Abstract

Deformation of the nucleus often presents a barrier to cell migration through tight spaces, such as those encountered as cells move through tissues or across extracellular matrix barriers. Reorganization of the nucleus to allow its passage through spaces much smaller than its resting diameter requires forces generated by the cytoskeleton, as well as active reorganization within the nucleus driven by ATPases that crosslink or move chromatin. Here, we show that two different nuclear ATPases, the BRG1/SMARCA4 motor of the BAF or SWI/SNF complex and the bifunctional crosslinking and loop extruding complex, cohesin, have opposite effects on the stiffness of isolated nuclei. Inhibition of BRG1 stiffens the nucleus, and cohesin softens it in karyoplasts derived from multiple cell types, including four different cancer cells, fibroblasts, and mesenchymal stem cells. The effects on isolated nuclear stiffness coincide with the effects of these ATPases on the ability of cells to migrate through tight spaces. Stiffening the nucleus inhibits single cell migration through micron-sized pores and the outward migration of tumor cell spheroids into a surrounding collagen matrix. Softening the nucleus by inhibiting cohesin has the opposite effect: it enhances single-cell migration through pores, at least for some cell types, and facilitates the outgrowth of cells from a tumor cell spheroid into the surrounding matrix. These results emphasize the importance of active motions generated within the nucleus for the global mechanics of the nucleus and the way that it deforms in response to externally generated stresses.

**Statement of significance:** Cells often move through tight spaces within three-dimensional materials. This constricted motion requires deformation of the nucleus, which is often stiffer and less dynamic than the rest of the cell. Nuclear deformation is achieved in part by forces generated outside the nucleus, primarily by cytoskeletal motors, but nuclear deformation is also affected by intranuclear motor proteins that move and reorganize chromatin. Here we show that the activity of DNA binding chromatin remodeling ATPases affects not only the stiffness of the isolated nucleus but also the ability of cells to deform as they move through tight spaces. Inhibition of motors such as BRG1 that fluidize the nucleus prevents cell movement through tight spaces, whereas inactivation of the crosslinking ATPase cohesin tends to enhance it.

## Introduction

The cell’s nucleus is often a limiting factor in the ability of cells to squeeze through tight spaces, such as those encountered during movement *in vivo* through tissues or *in vitro* through microfluidic channels or three-dimensional networks. Most models of nuclear deformation during cell motility focus on motor proteins and the active force generation by the cytoskeleton, which then transmits forces to the nucleus (1-4), typically modeled as a viscoelastic solid. Compliance of the nucleus to cell-generated forces can be altered by manipulating structures such as the nuclear lamina or the balance between euchromatin and heterochromatin, while the active motions are usually assigned to motor proteins such as myosin or kinesin outside the nucleus. Recently, the nuclear motor protein BRG1, a component of the SWI/SNF or BAF chromatin remodeling complex (1-3), was shown to have a large effect on the compliance of nuclei, isolated from fibroblasts as karyoplasts, to external forces (4). In this context, the BRG1 motor contributes to the active matter properties of the nucleus, enabling it to fluidize when subjected to external forces that cannot deform the nucleus to the same extent when this motor is inhibited, either pharmacologically or by depletion of ATP production. Whether other nuclear ATPases also alter nuclear compliance is not known, nor is the effect of inhibiting BRG1 on the mobility of intact cells through three-dimensional spaces. In this study, we show that inhibition of BRG1 significantly stiffens nuclei isolated as karyoplasts from a wide range of cells. In contrast, inhibition of the ATPase cohesin has the opposite effect, leading to softening of the nucleus for the same range of cell types. These opposite effects on nuclear deformation often lead to differences in the ability of intact cells to move either as single cells through micron-sized pores, or as cells escaping from a spheroid into the surrounding extracellular matrix. The effect of inhibiting these two motors is not the same in all cell types and appears to be blunted in some cell types, in which one or both of these nuclear ATPases are functionally altered (5,6). The results of this study show that nuclear ATPases that are mainly studied for their effects on transcription also modulate the effect of external forces, such as those generated by the cytoskeleton, to deform the nucleus to conform with the shape change of the whole cell. The large effects of inhibiting specific nuclear ATPases on cell and spheroid motility and outgrowth might have implications for the efficacy of pharmacological agents targeting these two motor proteins in therapeutic applications.

## Materials and methods

### Cell culture

Lung adenocarcinoma (A549), glioblastoma (LN18), breast cancer (MCF-7), hepatocellular carcinoma (Huh-7) and vimentin-null mouse embryonic fibroblasts (mEF -/-) cells were cultured in high-glucose DMEM (Corning, Corning, NY) supplemented with 10% FBS (Sigma-Aldrich, St. Louis, MO), 0.1 μg/mL penicillin, and 0.1 mg/mL streptomycin (Mediatech, Manassas, VA). Human mesenchymal stem cells (HMSCs) were grown in mesenchymal stem cell basal medium (440 mL) supplemented with growth supplement (50 mL), L-glutamine (10 mL), and GA-100 (0.5 mL) (all from Lonza, Walkersville, MD). Cells were maintained at 37□°C in a humidified 5% CO□ incubator and used for experiments between passages 3 and 10.

### Enucleation

Karyoplasts were isolated from fibroblasts, cancer, and mesenchymal stem cells as described previously (4) with minor modifications. Briefly, 100,000 cells were cultured on collagen-coated (0.1 mg/mL, overnight) glass cover slip inserts with a ring of polydimethylsiloxane (PDMS) on the edges for 24 h. Just before isolation, cells were incubated with 5 μg/mL Hoechst (diluted in cell culture medium) for 30 min. Subsequently, samples were centrifuged at 2600 × g for 50 min. The centrifugation was performed in DMEM supplemented with 1% FBS and 2 μg/mL cytochalasin D at 37°C using a Beckman-Coulter Optima LE-80 K ultracentrifuge with an SW-28 rotor. To stabilize the inserts during centrifugation, PDMS was cured in the base of ultracentrifuge tubes to create a flat surface. During centrifugation, isolated karyoplasts were collected in inverted inserts coated previously with 0.1 mg/mL poly-D-lysine (4°C, overnight). Enucleation efficiency was assessed by imaging cells after karyoplast isolation and analyzing the percentage of cells lacking nuclei, as determined by the absence of Hoechst staining. Karyoplasts were treated after isolation, for 2 hours, with a dual BRM and BRG1 inhibitor, BRM-014 (100 µM, iBRG1, Cayman Chemical, Ann Arbor, MI, #36138), a cohesin-inhibiting peptide with a TAT sequence (CIP3-TAT, Peptide 2.0, 10 µM), and doxorubicin (DOX, 100 µg/mL). An equal volume of solvent was added to the cells in vehicle control conditions. Following these treatments (after 2 h), karyoplasts were compressed using a Nanowizard 4 atomic force microscope (AFM, Bruker, Santa Barbara, CA), and then imaged after fixation using a Leica DMI8 confocal microscope (Leica, Wetzlar, Germany) with a 63× objective.

### AFM measurements

Atomic force microscopy (AFM) measurements were performed using a Bruker Nanowizard 4 system mounted on a Leica DMI6000 B microscope, as previously described (4). Karyoplasts were compressed using tipless silicon nitride cantilevers (NP-O10, Bruker) with a nominal spring constant of 0.35 N/m at a frequency of 1Hz, a speed of 20 µm/s, and a maximum applied force and strain of 50 nN and 30%, respectively. All measurements were made at room temperature in PBS, within 2-5 hours of karyoplast isolation. A minimum of 10 karyoplasts were measured per condition. Fluorescent and brightfield images were acquired for each karyoplast during AFM compression measurements to assess the relationship between karyoplast size and stiffness.

AFM force curves were analyzed using the JPK analysis software and fit to a double-contact hertz model for a sphere being compressed between two planes, as previously described (7), using the equation:

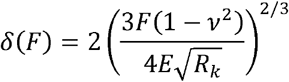

Where *δ* is the deformation as a function of the force *F, ν* is the Poisson’s ratio, *E* is the Young’s modulus and *R* _*k*_ is the radius of the karyoplast.

### Imaging

Karyoplasts and cytoplasts were fixed with 3.7% paraformaldehyde for 10□min at room temperature, then permeabilized with 0.1% Triton X-100 for 5□min. Samples were blocked in 0.1% bovine serum albumin (BSA) for 30□min. Isolated karyoplasts were incubated overnight at 4□°C with primary antibodies against lamin A/C (Cell Signaling Technology, 4777S) and lamin B (Abcam, ab16048), both diluted 1:500 in 1% BSA. Cytoplasts were incubated for 1□h at room temperature with phalloidin-DyLight 594 (Cell Signaling) and rabbit anti-vimentin antibody (1:500, Proteintech, 10366-1-AP), both diluted in 1% BSA. Following incubation with primary antibodies, samples were rinsed with 1% BSA and incubated for 1□h with secondary goat anti-mouse IgG with Alexa Fluor 488 (Invitrogen, A11029) and goat anti-rabbit IgG with Alexa Fluor 647 (Invitrogen, A-21244)—diluted 1:1000 in 1% BSA. After rinsing with PBS, samples were mounted using Anti-Fade Fluorescence Mounting Medium (Abcam) and imaged on a Leica DMI8 confocal microscope with a 63× objective.

### Transwell assay

To assess cancer cell transmigration through confined spaces, a Transwell assay (Corning) was performed using inserts with porous membranes featuring pores of 3, 5, or 8 µm in diameter (#3415, #3421, #3422). The lower chamber was filled with 500 μL of culture medium containing 5 ng/mL of transforming growth factor-beta 1 (TGF-β1, R&D Systems, Minneapolis, MN, #7754-BH/CF). In the upper chamber, 40,000 cells were seeded along with either DMSO, the BRG1 inhibitor (iBRG1; 25, 50, or 100 µM), CIP3TAT (10 µM), actinomycin D (100 µg/mL), or doxorubicin (100 µg/mL). After 4 h of incubation, non-migratory cells were removed from the upper membrane surface using a cotton swab. Migrated cells on the lower membrane surface were fixed in 3.7% paraformaldehyde for 10□min at room temperature and stained with Hoechst. Nuclei were imaged by fluorescence microscopy, and cell number and nuclear morphology were quantified using ImageJ.

### Migration assay

To assess cancer cell motility in response to BRG1 inhibition, cells were seeded onto collagen-coated 96-well plates at a density of 3,000 cells/well and allowed to adhere for 3 hours. Next (after 3 hours), BRG1 inhibitor (iBRG1, 100 µM), cohesin inhibitor (CIP3TAT, 10 µM), or DMSO (vehicle control) was added. All experiments were performed in the presence of 5 ng/mL TGF-β1 to mimic the Transwell assay setup. Time-lapse imaging was performed using a bright-field microscope equipped with an environmental chamber, capturing images every 10 minutes over a 24-hour period. Cell motility was quantified by tracking individual cells using ImageJ.

### Spheroids in collagen matrix

Spheroids were generated by seeding 3,000 cells in 100□µL of culture medium into each well of a 96-well ultra-low attachment plate (Corning, 7007). Plates were incubated at 37□°C in a CO□ incubator for 5 days to allow spheroid formation. After careful removal of the culture media, mature spheroids were embedded in 1 mg/mL or 2 mg/mL collagen type I hydrogels, prepared as previously described (8,9). Briefly, to prepare 100□µL of 1□mg/mL collagen solution, 10□µL of 10× PBS, 10□µL of 0.1□M NaOH, 70□µL of sterile distilled water, and 10□µL of 10□mg/mL collagen type I were mixed on ice.

For a 2□mg/mL collagen gel, the volume of collagen was doubled while maintaining the same total volume by reducing water content. The hydrogel mixture was gently added to wells containing spheroids and incubated at 37°C for 15 minutes to allow for polymerization. After gelation, 100 μL of culture medium containing DMSO, iBRG1 (100 µM), CIP3TAT (10 µM), or doxorubicin (100 µg/mL) was added to the top of the gel. Bright-field images (5× magnification) were taken every 24 hours for 3 days to quantitatively monitor spheroid morphology and outgrowth, which were later analyzed using ImageJ Fiji software. The area of each spheroid was first normalized to its own initial size at time 0, and then further normalized to the corresponding DMSO control. The data are therefore presented as fold change in spheroid area over time relative to DMSO.

### Cytotoxicity testing

Cytotoxicity of the tested compounds was assessed using the Live/Dead Viability Kit (Invitrogen, L3224). Mature spheroids were treated with iBRG1 (100 µM), CIP3TAT (10 µM), or doxorubicin (100 µg/mL) for 72□h, matching the duration of the collagen gel experiments. Calcein AM and ethidium homodimer-1 were then added according to the manufacturer’s instructions. After incubation, fluorescence images were acquired using a Leica DMIRE2 microscope to assess live (calcein-positive) and dead (ethidium-positive) cells.

### Genomic data source

Gene expression and functional dependency data were retrieved from the Cancer Dependency Map (https://depmap.org/portal) public release (25Q2). DepMap compiles data from the Cancer Cell Line Encyclopedia (CCLE) and large-scale CRISPR-Cas9 gene knockout screens. RNA expression values are reported as log□(TPM+1), (TPM – transcripts per million) while CRISPR dependency scores are calculated using the Chronos algorithm, which adjusts for copy number and experimental biases. RNA expression: log□(TPM+1) values >6 correspond to robust transcript abundance (∼TPM > 60), comparable to housekeeping gene levels. CRISPR dependency: A Chronos score of 0 indicates non-essentiality, whereas <0 corresponds to strong essentiality, with more negative values indicating stronger dependency. Genes that are both robustly expressed and display negative Chronos scores are generally considered biologically relevant and potentially critical for cell survival.

### Quantification and statistical analysis

Data are presented as median and□mean ±□SE from n independent experiments, as indicated in figure legends, unless otherwise stated. Statistical comparisons were performed using unpaired Student’s t-test (two groups) or one-way ANOVA followed by Tukey’s post hoc test (multiple groups). Differences were considered statistically significant at p□≤□0.05. Schematics were created with BioRender.

## Results

Karyoplasts, intact nuclei surrounded by a thin layer of cytoplasm and the plasma membrane, but no internal organelles or cytoskeleton, were prepared by centrifugation (4,10,11) from five different human cell lines, including bone marrow-derived mesenchymal stem cells (hMSC), A549 lung cancer cells, LN18 glioma cells, MCF7 breast cancer cells, and Huh7 liver cancer cells (Figure 1A,B).

**Figure 1.**
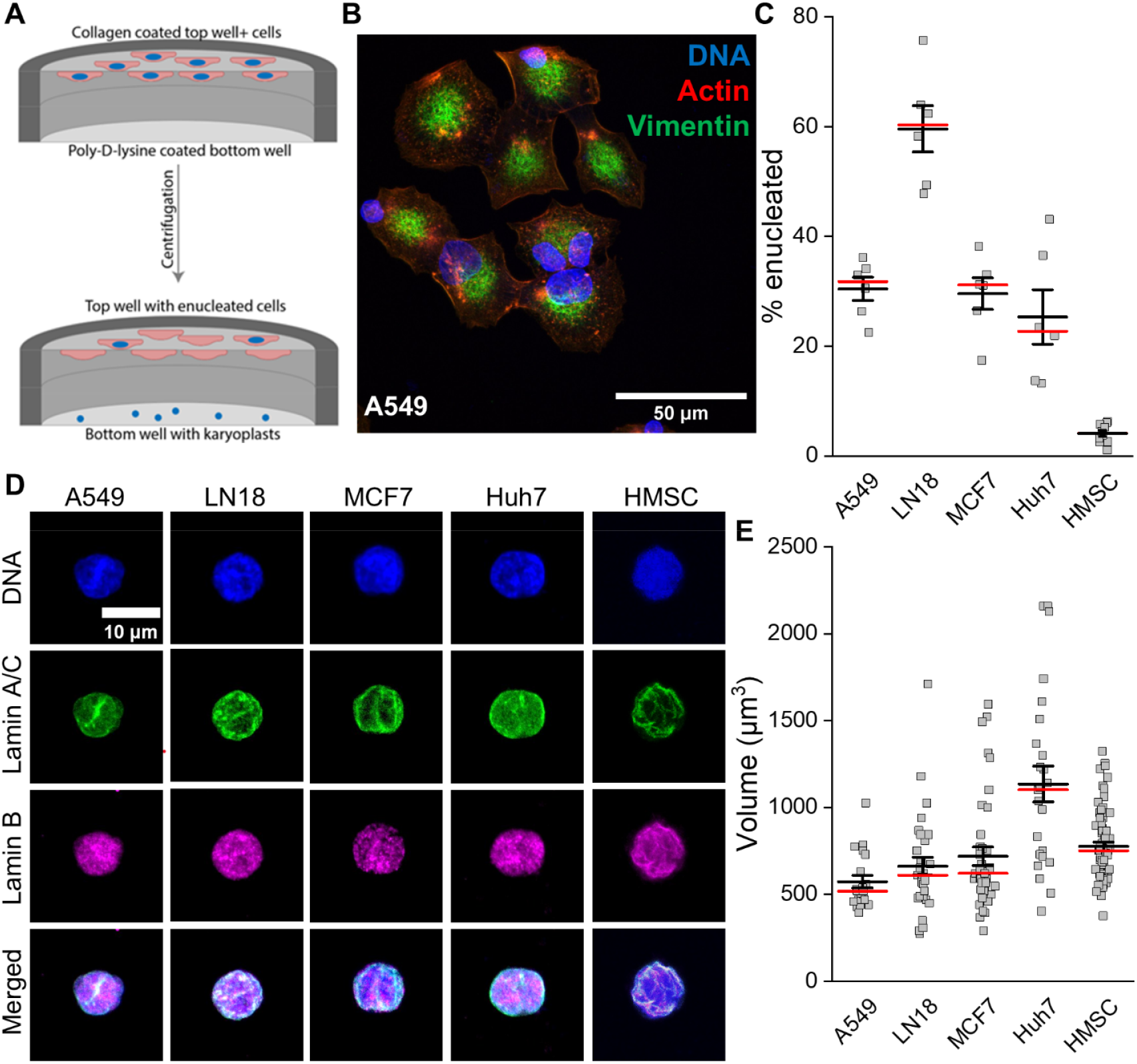
Generation and characterization of enucleated cells and isolated nuclei. (**A**) Schematic representation of the enucleation process. (**B**) Representative fluorescence image of enucleated A549 cells, highlighting the presence of cytoplasts lacking nuclear material. Cells are stained for vimentin (green), actin (red), and DNA (blue). Scale bar, ∼50 µm. (**C**) Quantification of enucleation efficiency, expressed as the percentage of cytoplasts relative to the total cell population. **(D)** Fluorescence images of isolated nuclei stained for DNA (blue), anti-lamin A/C (green), and anti-lamin B (magenta). Scale bar, ∼10 µm. (**E**) Mean volume of isolated nuclei. Data are presented as median (red) and mean ± SE; n ≥ 3 of independent experiments.

Karyoplasts could be isolated with variable efficiency from each of these cell types, as shown in Figure 1C. The appearance of the nuclei was determined by staining for chromatin within the nucleus interior and lamins A/C and B to demarcate the nuclear envelope, as shown in Figure 1D. Consistent with previous results from karyoplasts isolated from fibroblasts (4), these nuclei are approximately spherical with condensed chromatin concentrated in a subfraction of the nuclear interior and a folded nuclear lamina visualized by lamin A/C staining, with lamin B forming variable but more punctate patches on the surface or within the nucleus. Just as with whole cell volume, the volume of nuclei within karyoplasts also shows a wide variance (12), but with average values significantly different for each cell type (Figure 1E).

Nuclear stiffness, measured by whole karyoplast uniaxial compression (Figure 2A,B) as quantified by an apparent Young’s modulus (Figure 2C), also varied widely for karyoplasts isolated from different cell types, as did the degree of dissipation (Figure 2D) during a cycle of deformation and recovery of the nucleus. Nuclei isolated from the breast cancer cell line MCF7 and the hepatocellular carcinoma cell line Huh7 were the softest among the cell types compared, with the nuclei from lung cancer and glioma cells approximately 2 times stiffer, and close to the elastic modulus of karyoplasts isolated from fibroblasts. The mechanical dissipation during a cycle of deformation or recovery, which is a measure of the amount of energy dissipated compared to the energy elastically stored when the nucleus is compressed, also varied widely among different cell types. MCF7 and Huh7 cells not only had the lowest elastic modulus but also exhibited the highest relative amount of dissipation compared to the other three cell types.

**Figure 2.**
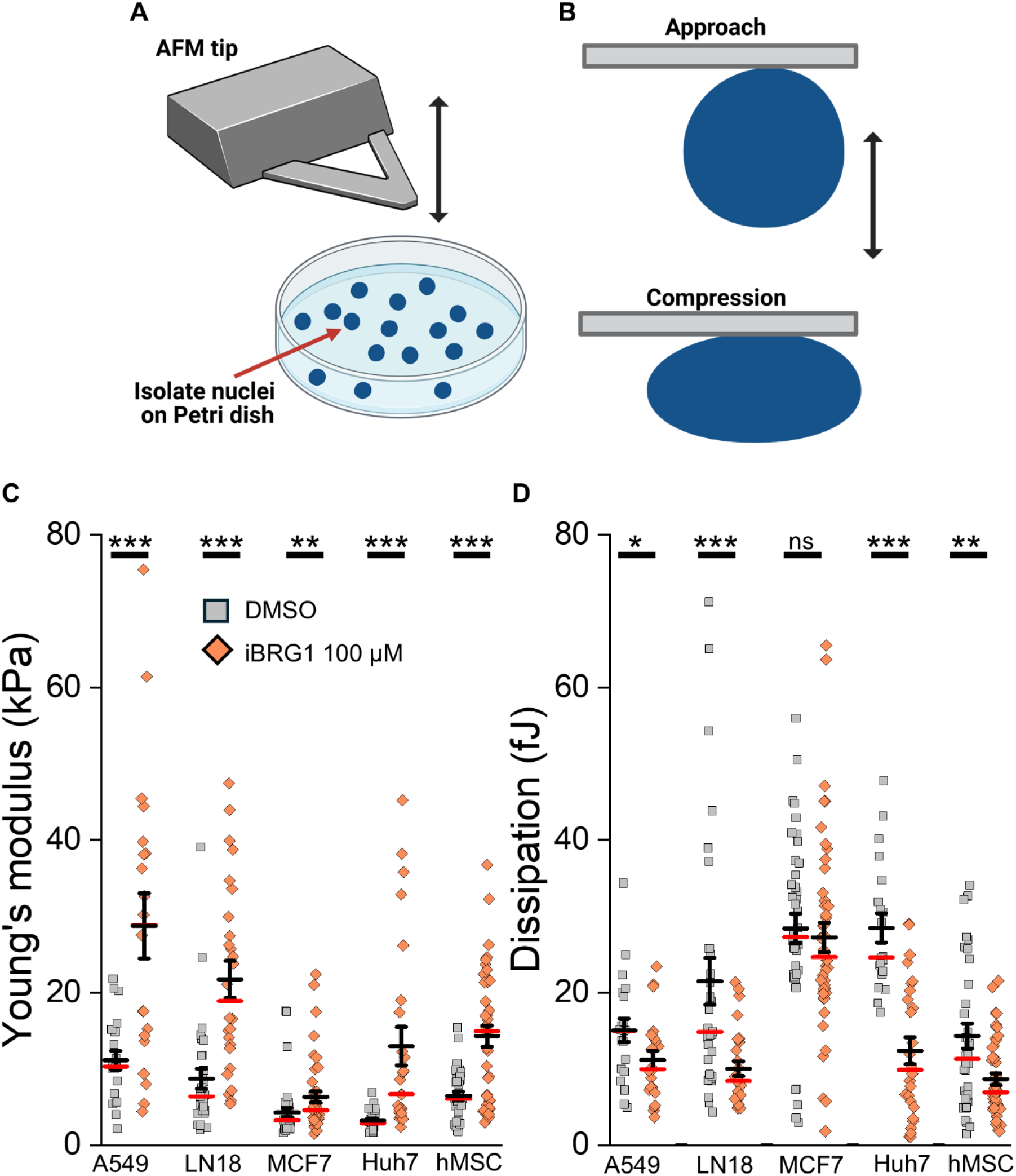
Effect of BRG1 inhibition on karyoplast deformation. (**A, B**) Schematic of the geometry of karyoplast deformation by a flat AFM cantilever. Effect of BRG1 (100 µM) inhibition on the apparent Young’s modulus (**C**) and dissipation (**D**) during a cycle of compression and recovery. Data are presented as median (red) and mean ± SE; n ≥ 3 of independent experiments. *, P□≤□0.05; **, P□<□0.01; ***, P□<□0.001. Significance was determined by an unpaired Student’s t-test.

As seen previously with karyoplasts isolated from fibroblasts, both the elastic modulus and the degree of dissipation were altered after the BRG1 chromatin remodeling motor was inhibited pharmacologically, but with quantitatively distinct effects on different cell types. BRG1 inhibition led to an approximately three fold increase in Young’s modulus of fibroblast nuclei, and a similar decrease in the degree of dissipation (4). A similarly large increase in Young’s modulus was observed when BRG1 was inhibited in A549, LN18, or hMSC cells, with a smaller effect on MCF7 and Huh7 cells. The decrease in mechanical dissipation was also variable among cell lines, with large effects on LN18 and Huh7 cells but smaller effects on A549 and MCF7 cells.

In contrast to the effect of inhibiting BRG1, which makes karyoplasts stiffer and less dissipative, inhibition of the DNA loop-regulating protein cohesin had opposite effects on karyoplast stiffness in all cell types, as shown in Figure 3. Cohesin inhibition with CIP3TAT decreased the apparent Young’s modulus of all six cell types (Figure 3A), but unlike BRG1 inhibition, it increased dissipation in some of the cell types (Figure 3B). The largest effects of cohesin inhibition on both Young’s modulus and dissipation were seen in LN18 cells. We also evaluated the effect of the topoisomerase inhibitor, doxorubicin, but we did not observe changes in the stiffness of the cells (Supplemental Figure 1).

**Figure 3.**
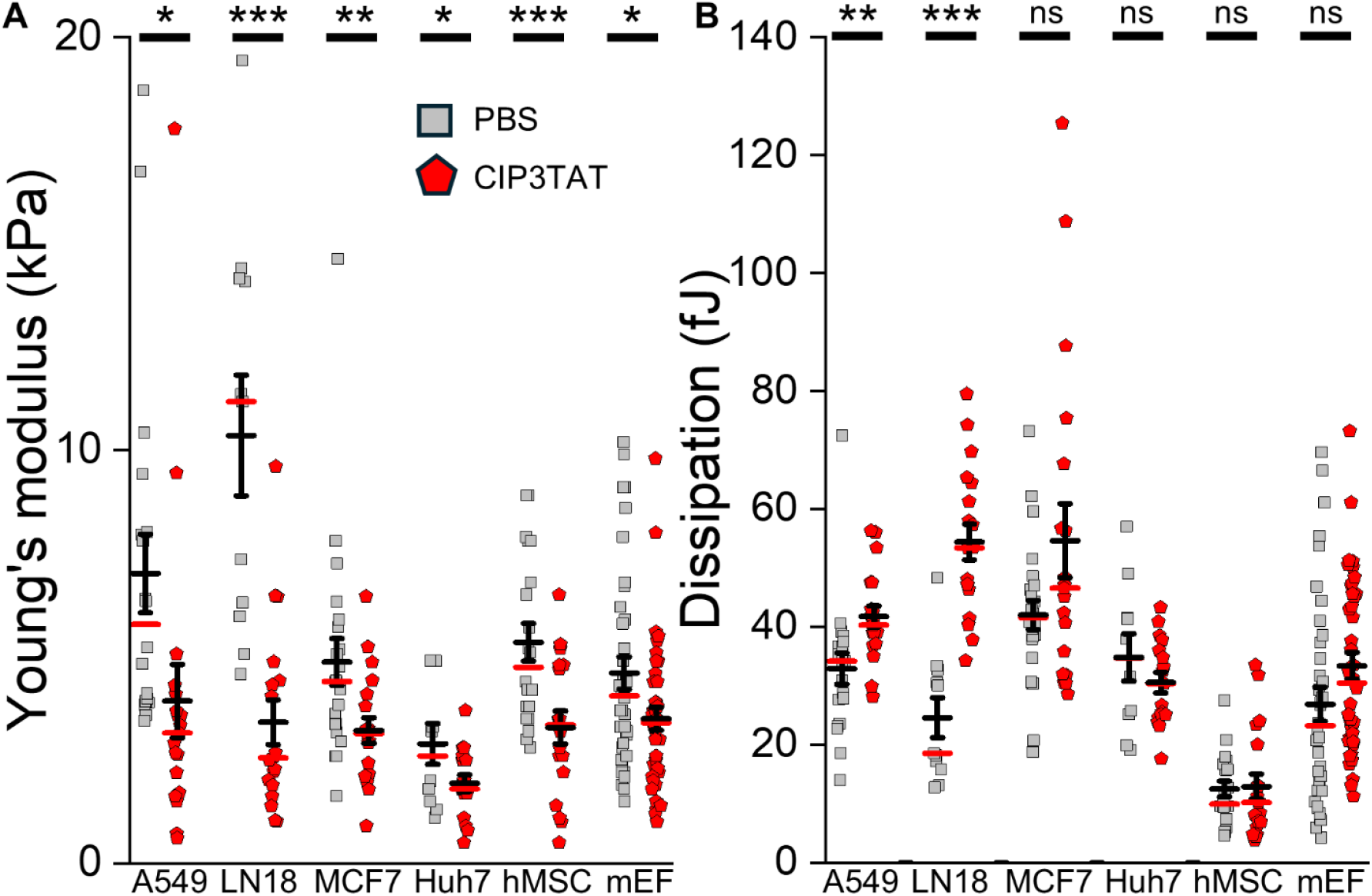
Effect of cohesin inhibition (CIP3TAT, 10 µM) on the apparent Young’s modulus (**A**) and dissipation (**B**) during a cycle of compression and recovery. Data are presented as median (red) and mean ± SE; n ≥ 3 of independent experiments. *, P□≤□0.05; **, P□<□0.01; ***, P□<□0.001. Significance was determined by an unpaired Student’s t-test.

The effect of inhibiting these two nuclear ATPases on nuclear stiffness has been primarily studied by application of inhibitors to isolated karyoplasts, but previous studies have also shown that addition of the BRG1 inhibitor to intact cells strongly diminishes the efficiency with which nuclei can be removed from cells for the centrifugation method required to produce karyoplasts (4), suggesting that in the impacted cell, BRG1 inhibition also increases nuclear stiffness, preventing its shape change required as the oblate ellipsoid nucleus of a well-adhered cell transforms into the spherical nucleus that emerges in the karyoplasts. These results motivated studies to test whether pharmacologic inactivation of nuclear ATPases would affect cell migration through tight spaces that require large deformations of the nucleus. These experiments were done either in single cells passing through micron sized pores, or with tumor cells spheroids embedded in a 3D collagen network.

As a control to determine if inhibiting BRG1 and cohesin altered the cytoskeletal machinery driving motility on a time scale of hours after addition of the inhibitors, unconstrained motility in 2D on the surface of tissue culture dishes was first measured. BRG1 inhibition had no statistically significant effect on migration rates after 4 h in any cell line. After 24 h, LN18 and Huh7 cells remained unaffected, whereas A549 and MCF7 cells showed moderate reductions (<30% and <10%, respectively; Figure 4). Cohesin inhibition produced a similar effect on A549 cells at 24 h but also slowed LN18 migration at both 4 h and 24 h, with no measurable effect on MCF7 or Huh7 cells.

**Figure 4.**
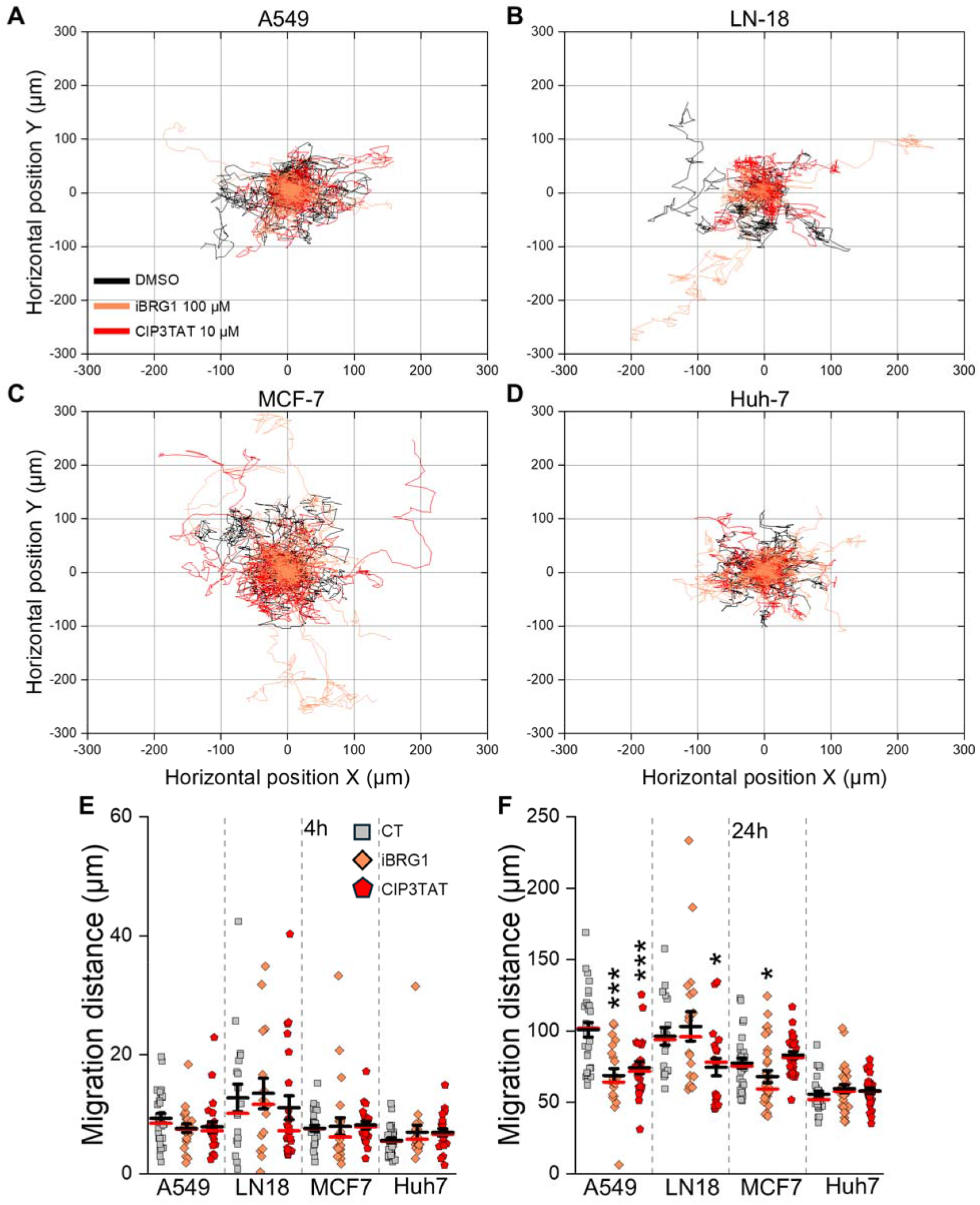
Inhibiting BRG1 and cohesin has a negligible acute effect on 2D cell motility. Tracks of cell centroids during motility of 4 cancer cell lines with or without inhibition of BRG1 (100 µM) and cohesin (CIP3TAT, 10 µM) (**A-D**) and total migration distances after 4 hours and 24 hours (**F**). Data are presented as median (red) and mean ± SE. At least 30 individual cells were tracked in each condition. *, P□≤□0.05; ***, P□<□0.001. Significance was determined by an ANOVA with post-hoc Tukey’s test.

To test cell migration in constricted spaces, pharmacological agents that inhibited the activity of BRG1 (iBRG1), cohesin (CIP3TAT), topoisomerase (doxorubicin), and RNA polymerase (Actinomycin D) were added to cells immediately before their addition to a transwell migration assay system in which cells chemotax in a growth factor gradient through pores of variable sizes (Figure 5A,B).

**Figure 5.**
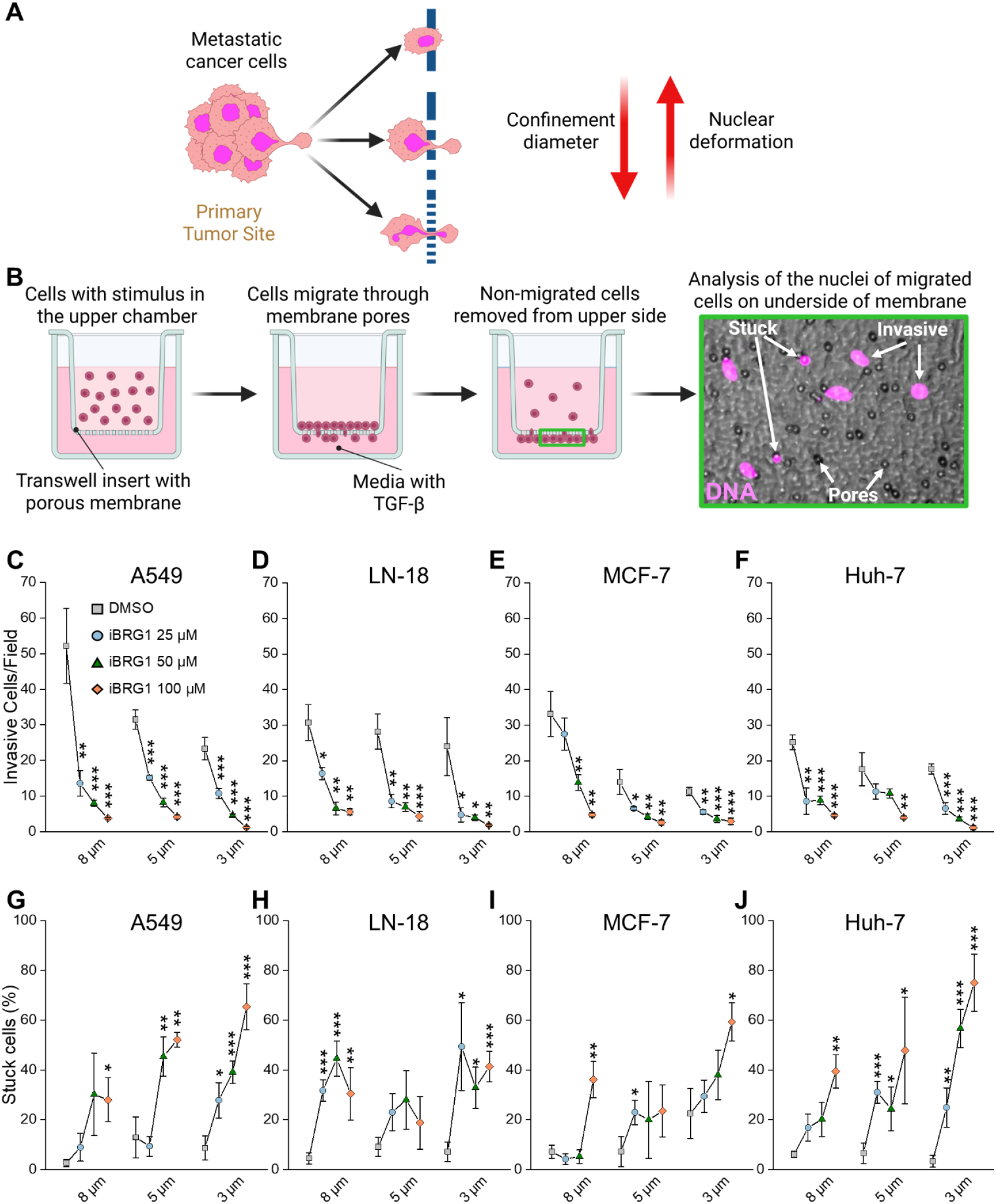
Effect of inhibiting BRG1 on the migration of cancer cells through micron-sized pores. (**A**,**B**) Schematic diagrams of constricted migration during metastasis and *in vitro* through membrane pores. Dose-dependent effect of BRG1 (25, 50, and 100 µM) inhibitor on migration of A549 (C), LN18 (D), MCF7 (E), and Huh7 (F) cells through pores of different sizes after 4 hrs. Parallel analysis of the fraction of cells trapped within the pores after 4 hrs (**G-J**). Data are presented as mean ± SE; n ≥ 4 of independent experiments (5-10 images each). *, P□≤□0.05; **, P□<□0.01; ***, P□<□0.001. Significance was determined by an unpaired Student’s t-test, comparing iBRG1-treated cells vs. DMSO-treated cells within each pore size condition.

Doxorubicin and Actinomycin D were used as a control, since they are ATPases, just as BRG1 and cohesin, but do not affect the stiffness of the nucleus, as seen in Supplemental Figure 1 and our previous report (4). Figure 5C-F shows that the BRG1 inhibitor decreases the ability of all four cancer cell types to migrate through pores in a dose-dependent manner. As the pore size became smaller, from 8 microns to 3 microns, the effect of BRG1 inhibition increased. Coincident with the decreased number of cells capable of migrating through the pore, there was an increase in the fraction of cells that became trapped within the pores as the BRG1 inhibitor concentration was increased (Figure 5G-J). This result suggests that BRG1 inhibition does not globally prevent cell motility, but rather increases the probability that a cell will migrate into a pore yet will not be able to traverse it within the four hours over which pore migration was measured. Additionally, we conducted parallel experiments simulating what occurs on the top of the insert membrane (Supplemental Figure 2). It shows that cells treated with different ATPases do not significantly differ from untreated cells in terms of morphology and their ability to adhere and spread after four hours of stimulation.

Even for cells that were able to pass through pores, the effect of BRG1 inhibition on nuclear mechanics was evident from the apparent shape changes in the nucleus seen in cells after they passed through the tight constriction, especially for the smallest pore diameter, as illustrated in Figure 6. After a control cell passes through a pore that requires nuclear deformation, the nucleus generally recovers its original shape without damage to the nuclear membrane, as reflected by a projected area that is near that of the same cell type on a flat surface. Depending on the cell type, addition of the BRG1 inhibitor shows a small but significant increase in the nuclear area and aspect ratio after a cell has traversed a pore with a diameter of eight or five microns, and a much larger effect on cells passed through three-micron pores (Figure 6A-H). This large, unrecovered shape change is illustrated for all four cell types in the images in Figure 6I, which show large, irregular Hoechst-stained chromatin, likely reflecting a broken nuclear membrane, given the smears of DNA outside the nuclear border. A similar effect was observed for fibroblasts (Supplemental Figure 3A). For HMSCs, the trend was also maintained; these cells were less likely to migrate, with most remaining trapped, even in relatively large pores (8 µm), likely due to their large nuclear surface area (Supplemental Figure 3B-D).

**Figure 6.**
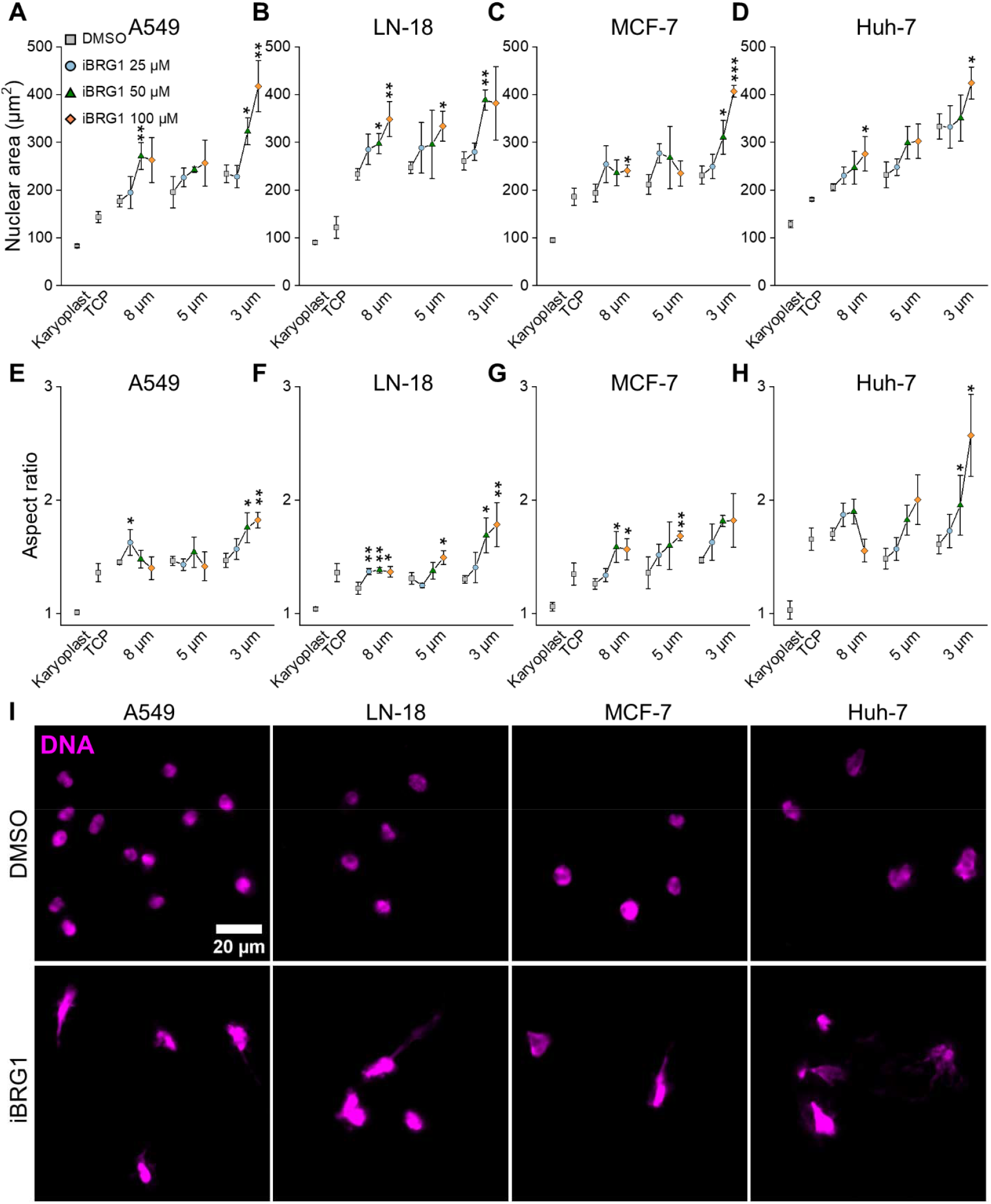
Effect of BRG1 inhibition on nuclear morphology in cells after migration through pores. The projected nuclear area (**A-D**) and aspect ratio (**E-H**) in cells that have passed through pores of various sizes, compared to the area of nuclei on flat surfaces before passage and within karyoplasts. (**I**) Images of nuclei after passage of cells through 3-micron pores (iBRG1 at 100 µM). Data are presented as mean ± SE; n ≥ 4 of independent experiments (5-10 images each). *, P□≤□0.05; **, P□<□0.01; ***, P□<□0.001. Significance was determined by an unpaired Student’s t-test, comparing iBRG1-treated cells vs. DMSO-treated cells within each pore size condition.

In contrast to the strong effect of BRG1 inhibition in decreasing migration through pores observed in all four cancer cell types, inhibiting cohesin with a cell-permanent inhibitor significantly increased the motility of three of the four cancer cell types (Figure 7). A549 and Huh7 cells were particularly strongly affected by cohesin inhibition, with large increases in the number of cells passing through 3-micron pores. In contrast, LN18 cells were not enhanced to pass through small pores after cohesin inhibition, but rather showed a tendency to decrease migration. The opposite effect of CIP3TAT on LN18 migration could be attributed to their robust RNA expression and negative CRISPR score (Supplemental Figure 4). Inhibition of RNA polymerase by actinomycin D had no systematic effect on cell migration through pores, as shown in Figure 7. Inhibition of topoisomerase by doxorubicin slightly decreased migration through pores, presumably because doxorubicin, unlike BRG1 inhibitors, is cytotoxic, as evidenced by previous reports at similar doses and times (13).

**Figure 7.**
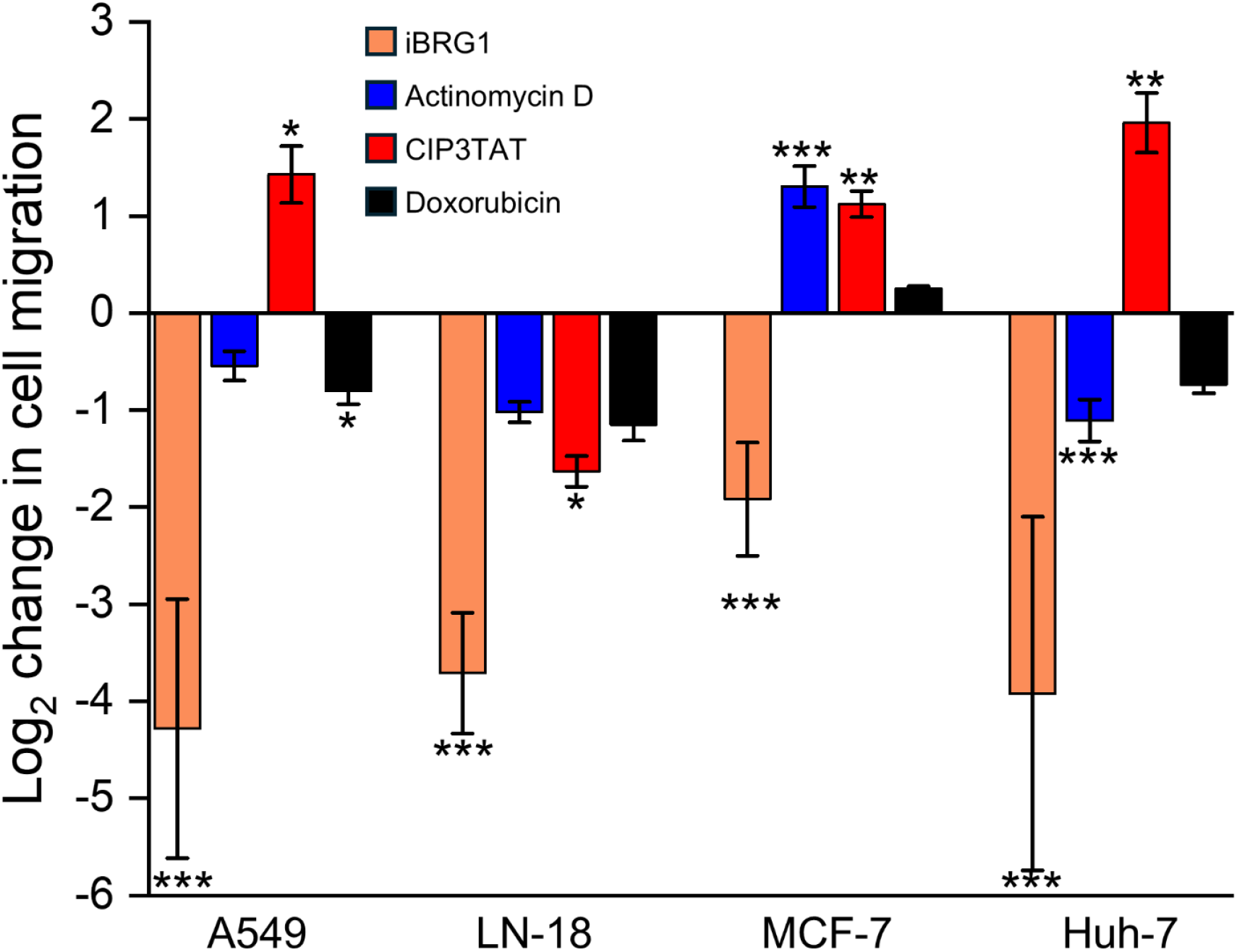
Effect of inhibiting BRG1 (100 µM), Actinomycin D (RNA polymerase, 100 µg/mL), cohesion (CIP3TAT, 10 µM), and doxorubicin (topoisomerase, 100 µg/mL) on cell motility through 3 micr n pores. Data are presented as mean ± SE; n ≥ 3 of independent experiments. *, P□≤□0.05; **, P□<□0.01; ***, P□<□0.001. Significance was determined by an unpaired Student’s t-test, compared to the untreated condition.

The relevance of altering nuclear mechanics for cells migrating in three-dimensional matrices was studied by placing tumor cell spheroids within the collagen networks and then monitoring the outgrowth of cells from the spheroid perimeter into the surrounding matrix. Figure 8 shows a control experiment in which spheroids were formed from LN18 and A549 cells in suspension and then treated with the four nuclear ATPase inhibitors for 72 hours, the time over which migration into the matrix was measured. MCF7 and Huh7 cells did not form spheroids under these conditions. These images demonstrate that inhibition of BRG1 and cohesin has a negligible effect on the integrity of the tumor spheroids in suspension, with no significant evidence of cell death resulting from these inhibitors. In contrast, doxorubicin treatment resulted in a significant increase in cell death, as quantified by a live-dead assay, consistent with the effective inhibition of cell migration observed in Figure 7, which is attributed to cell toxicity rather than a defect in nuclear deformation.

**Figure 8.**
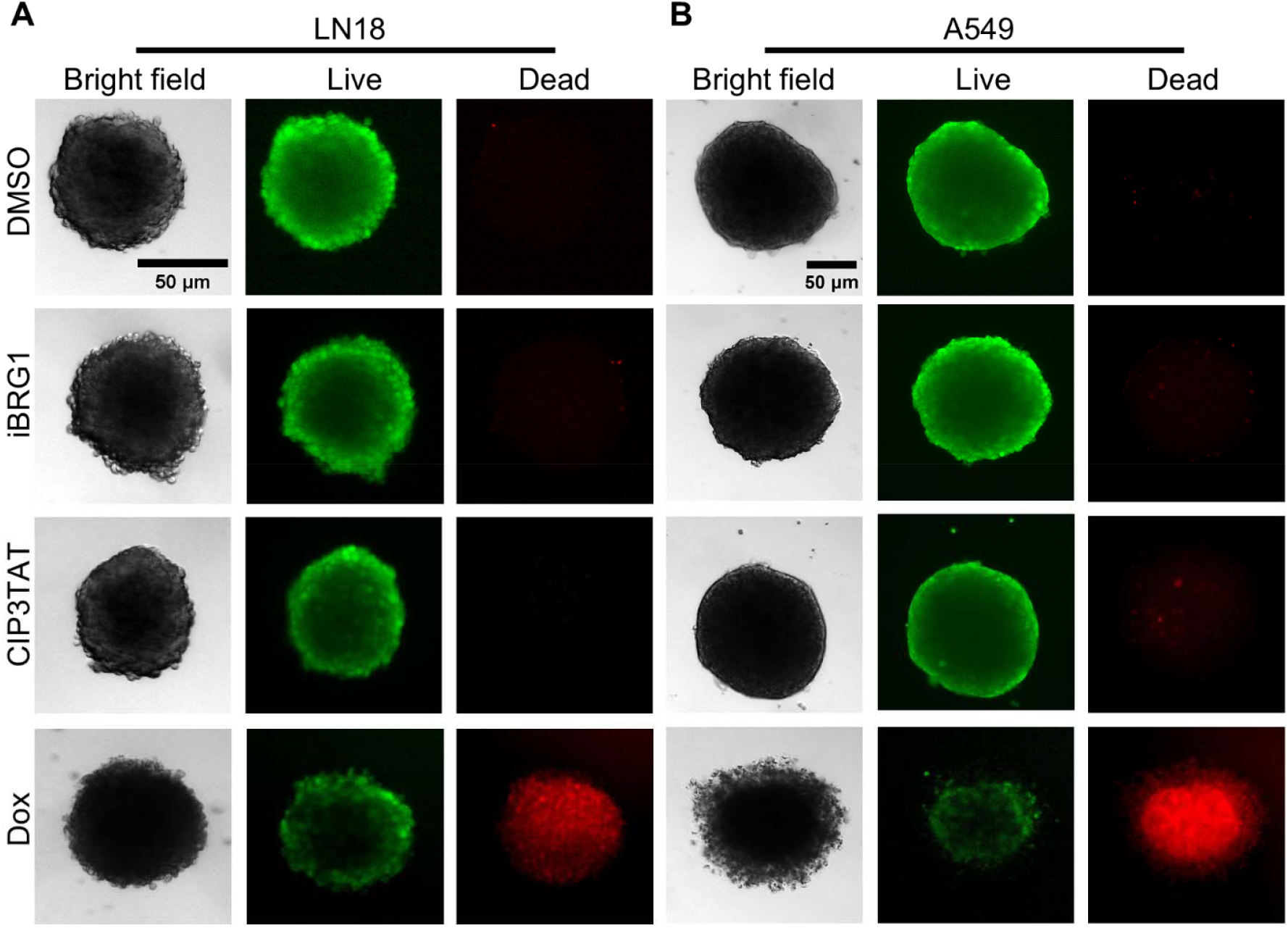
Viability of spheroids after inhibition of BRG1 (iBRG1, 100 µM), cohesin (CIP3TAT, 10 µM), and topoisomerase (doxorubicin, 100 µg/mL). Live/dead stains of LN18 (**A**) or A459 (**B**) spheroids 72 hours after adding inhibitors in suspension culture. n ≥ 3 independent experiments.

When spheroids such as those shown in Figure 8 are placed into a collagen network, they no longer always retain their spherical shape, but, depending on the chemical and physical features of the matrix surrounding them and the strength of cell-cell adhesions, cells at the spheroid boundary can migrate into the matrix and invade it. Figure 9 shows how spheroids of LN18 glioma cells diffusely migrate in the collagen networks, with increased migration efficiency as the collagen concentration increases and the mesh size of the network decreases (∼8 µm for 1 mg/mL and ∼2 µm in 2 mg/mL), presumably because of both increased integrin activation after attachment to collagen and the ability to generate larger traction stresses at the spheroid-matrix interface (14). Addition of the BRG1 inhibitor largely prevents this outward migration of cells, with the effect being greater for the more concentrated network (Figure 9A), where the smaller mesh size and stiffer gel would require a greater deformation of the nucleus. The time course of spheroid outgrowth, shown in Figure 9B,C, confirms that spheroid growth is decreased by BRG1 inhibition in a dose-dependent manner.

**Figure 9.**
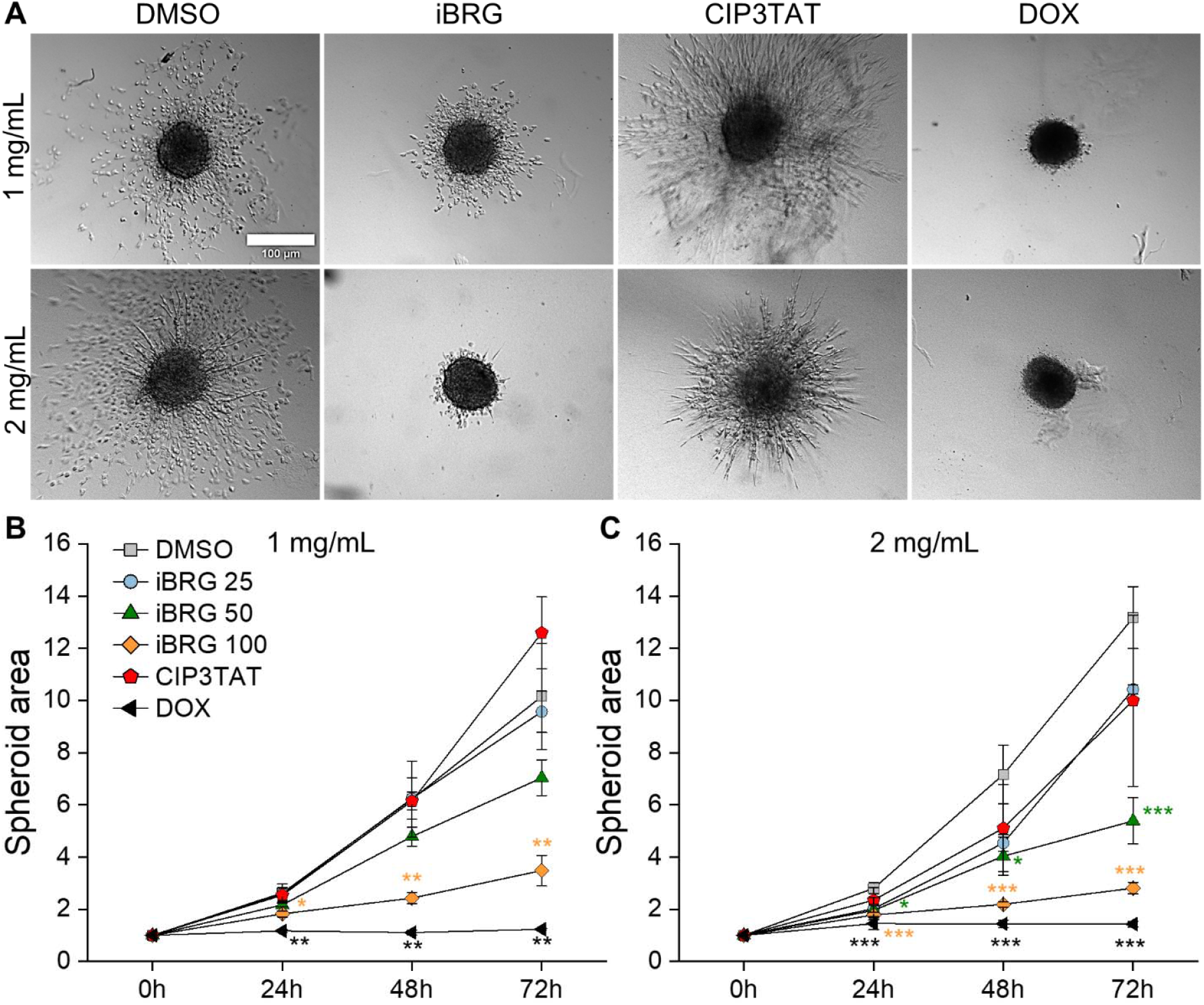
Effects of nuclear ATPases on LN18 spheroid migration into collagen matrices. Outgrowth from spheroids 72 hr after encapsulation in 1 (**A**) or 2 (**B**) mg/ml collagen gels. Total spheroid area after addition of inhibitors of BRG1 (iBRG1, 25-100 µM), cohesin (CIP3TAT, 10 µM), or doxorubicin (100 µg/mL) (**C, D**). n ≥ 3 independent experiments, at least 3 spheroids per experiment. Data are presented as the mean□±□SE. *, P□≤□0.05; **, P□<□0.01; ***, P□<□0.001. Significance was determined by one-way ANOVA with Tukey’s test, compared to the DMSO condition.

Doxorubicin completely suppresses spheroid outgrowth, as expected from its cytotoxic effect, but cohesin inhibition does not decrease spheroid outgrowth, and possibly increases it for the lower concentration collagen gel. Cohesin inhibition has an impact, possibly to increase the number of cells that migrate out of the spheroid, but also to transform the outward migration from that of single cells migrating into the matrix for the controls to streaks or sheets of cells that migrate into the matrix while maintaining cell-cell contact.

A similar effect of BRG1 inhibition in preventing tumor cell outgrowth from A549 lung cancer spheroids is shown in Figure 10A. A549 spheroids are much larger than those formed by LN18 cells, and their outgrowth into the matrix is somewhat more dependent on the concentration and, therefore, the mesh size of the collagen network. Similar to the effect seen in Figure 9, BRG1 Inhibition strongly suppresses migration into the matrix, but inhibition of cohesin does not do so as strongly (Figure 10B,C). Doxorubicin also prevents outward migration, again presumably due to its cytotoxic effects.

**Figure 10.**
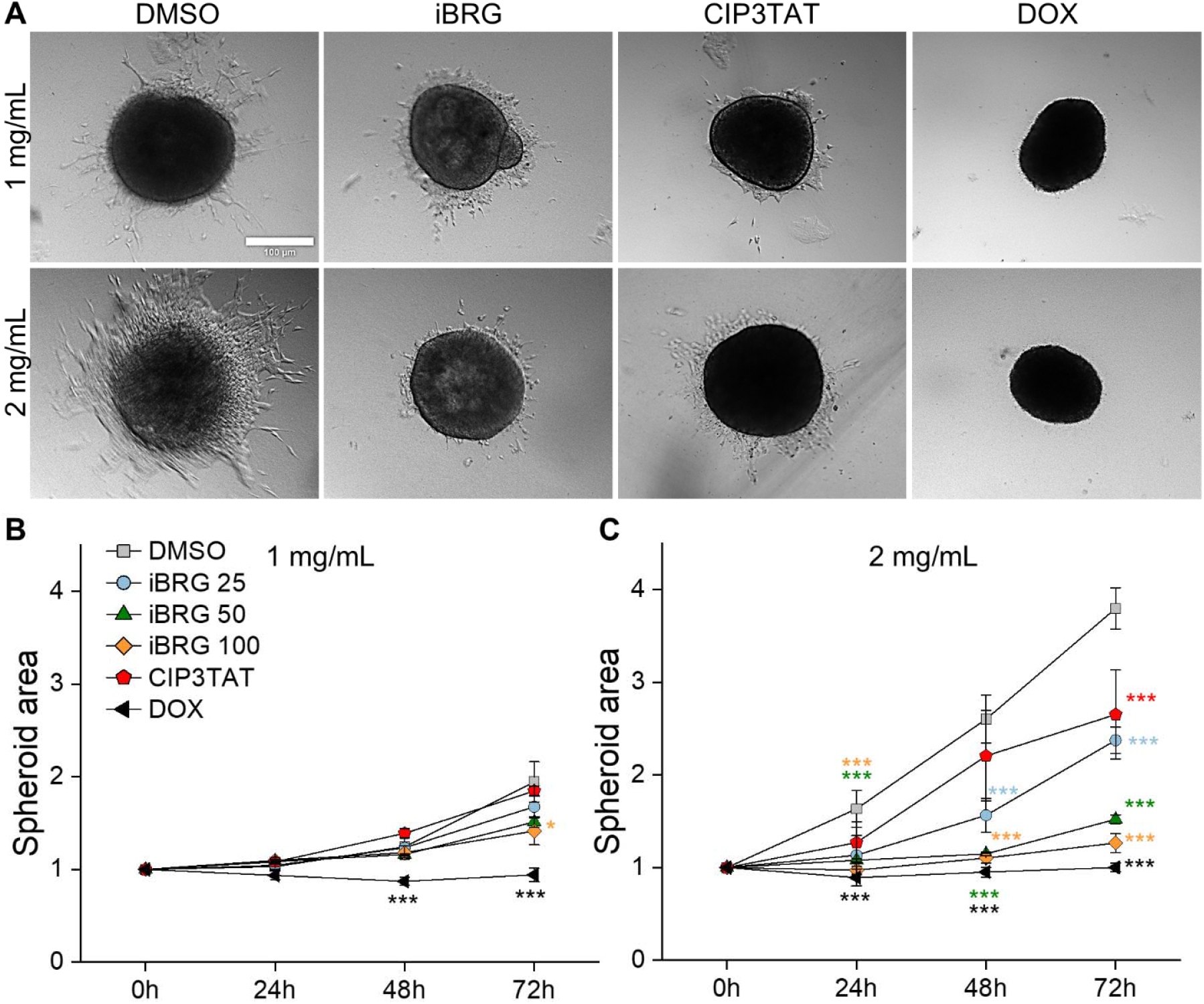
Effects of nuclear ATPases on A549 spheroid migration into collagen matrices. Morphology and outgrowth from spheroids 72 hr after encapsulation in 1 (**A**) or 2 (**B**) mg/ml collagen gels. Total spheroid area after addition of inhibitors of BRG1 (iBRG1, 25-100 µM), cohesin (CIP3TAT, 10 µM), or topoisomerase (doxorubicin, 100 µg/mL)(**C, D**). n ≥ 3 independent experiments, at least 3 spheroids per experiment. Data are presented as the mean□±□SE. *, P□≤□0.05; **, P□<□0.01; ***, P□<□0.001. Significance was determined by one-way ANOVA with Tukey’s test, compared to the DMSO condition.

## Discussion

Deformation of the nucleus is generally not an essential aspect of cell motility *in vitro* on flat surfaces, but becomes important and sometimes rate-limiting when cells need to squeeze through the tight spaces they encounter *in vivo* within tissues. In the context of cancer, massive deformation of the nucleus is required when cells breach the extracellular matrix and invade neighboring tissues, where the spaces between matrix fibers are often much smaller than the typical diameter of the nucleus, which is in the range of 10 microns, as shown in Figure 1. The structures that determine the mechanical resistance of the nucleus to deformation and the forces that are generated to deform it are only partly understood and likely to involve multiple different structures and force-generating ATPases. Forces generated outside the nucleus are essential for global cell motility as well as for applying mechanical stresses to the nuclear surface, and therefore, treatments such as inhibiting cytoskeletal myosin motors have shown efficacy in limiting the outgrowth of rapidly invasive tumors such as gliomas (15,16).

If the nucleus is, as often assumed, the stiffest organelle in the cell, then large deformation of the nucleus requires greater mechanical stress than the elastic modulus of the nucleus, often estimated to be in the range of a few kPa. However, nuclei undergo large deformations when, for example, lipid particles impinge on them within a fat-filled hepatocyte (17,18), even though the contractile force of the hepatocyte applies only a few hundred Pascal stress to the nuclear surface (17). For biological materials, stiffness cannot be quantified by a scalar quantity, and the elastic modulus, as well as the dissipation within the material, depend on the magnitude and duration of deformation. Numerous studies have demonstrated that the nucleus exhibits complex rheological responses, including power-law viscoelastic relaxation (19) and time-dependent deformability, which at long times leads to liquid-like flow (19). These rheological responses are a combination of both the viscoelastic properties of the chromatin and other macromolecular components crowded within the nucleus, the elastic properties of the nuclear lamina composed of both the protein lamin network and the phospholipid double bilayer membrane, but also the active processes that drive these structures out of equilibrium and create an active composite material. Numerous studies have shown that active, ATP-driven random motions, whether in the form of actomyosin motor movements (20) or the shape fluctuations of ATP-dependent enzymes (19), can fluidize materials within the cytoplasm of both eukaryotic and prokaryotic cells. Similar active matter properties are also expected to be prominent in the nucleus, which contains abundant ATPases involved in chromatin reorganization, elongation of DNA and RNA chains, breakage and reformation of DNA double strands, and other active processes. Both BRG1/BRM and cohesin are relatively abundant nuclear proteins. Previous studies in yeast report ∼200 and 2000 of the related SWI/SNF and RSC complexes, respectively, each of which has a BRG1/BRM-like motor (21). This corresponds to a concentration of 0.9 µM, assuming a 2 µm diameter yeast nucleus. Cohesin protein is expressed at 250,000 copies in a 10 µm diameter mammalian nucleus, corresponding to 0.8 µM (22). The expression level of these enzymes depends on cell type and condition, so these are only a rough guide. The RNA expression levels and CRISPR scores for the BRG1 and cohesin complexes in all tested cancer cell lines are shown in Supplemental Figure 4.

Previous studies have shown that inhibition of the BRG1/SMARCA4 motor, the active component of the chromatin remodeling BAF or SWI/SNF complex, increases the stiffness of an isolated nucleus within a karyoplast and prevents the release of flattened nuclei within well-adherent cells in response to centrifugal forces that otherwise are sufficient to form karyoplasts from control cells (4). The initial studies were done with mouse embryo fibroblasts, and here we show that similar effects of nuclear stiffening and loss of dissipation are observed in nuclei isolated from mesenchymal stem cells, as well as from four different cancer cell types. The size of the nuclei isolated from these different cells, as well as their apparent elastic modulus and the degree of dissipation as the whole nucleus is deformed, vary from cell type to cell type, but in all cases, inhibition of BRG1 causes the nuclei to become stiffer and less dissipative.

These studies also show that the effects of nuclear stiffening and the loss of energy dissipation when the nucleus deforms, caused by inhibition of BRG1, lead to significant differences in the ability of intact cells to move through tight spaces, either as single cells through micron-sized pores, or as individuals or cell collectives leaving a spheroid embedded within a collagen matrix. Dynamic remodeling of chromatin is essential for large deformations of the nucleus, and to the extent that these dynamics depend on the actin motions of the BAF complex or other chromatin remodeling motors, their inhibition will tend to slow the rate or limit the extent to which the nucleus deforms. Since the BAF complex, powered by BRG1 motion, is essential for the reorganization of chromatin to allow for the exposure of DNA required for transcription of specific genes, inhibition of this motor undoubtedly leads to protein expression changes that are likely to affect cell viability or motility.

However, single-cell migration through porous filters requires only a few hours, strongly supporting the hypothesis that the effect of inhibiting BRG1 on the decreased ability to migrate through tight spaces results from acute changes in nuclear stiffness that then lead to increased numbers of cells trapped within the pores through which the cells are attempting to crawl. The longer times required to grow tumor cell spheroids and monitor their outward migration through collagen gels prevent a straightforward interpretation of the effects of BRG1 inhibition on the outward migration of tumor cells from a spheroid into a surrounding matrix, as shown in Figures 9-11. However, the finding that the effect of inhibiting BRG1 on outward migration from the spheroid is more substantial in denser collagen gels is also consistent with the idea that a stiffened nucleus, due to the loss of this motor activity, is an essential element in the inability of cells to migrate out from the spheroid. Control experiments show no effect of BRG1 inhibition on cell viability in either of these motility assays, again supporting the idea that mechanical effects, rather than other defects within the cell, account for their inability to move through tight spaces.

The effect of inhibiting BRG1 on cell migration might be relevant to interpreting the effects of pharmacological use of BRG1 inhibitors on tumor growth *in vivo*. Elements of the BAF complex are mutated or their expression levels are altered in a significant fraction of human cancers, with abnormal expression of BRG1 particularly clearly documented in some forms of gliomas (23-27) as well as hepatocellular carcinoma (28) and prostate cancer (29). Previous studies have shown that BRG1 overexpression is highly correlated with poor prognosis (25,29), and that pharmacologic inhibition of BRG1 might lead to decreased tumor growth (26,27,29). Whether these effects depend entirely on the role of BRG1 in transcription and cell proliferation, or whether some of the beneficial effects derive from the inability of tumor cells to move into the tight spaces of the microenvironment is not yet clear. Previous studies have shown that BRG1 is upregulated in MCF7 cells and that downregulating its expression with siRNA decreases migration (6). Silencing BRG1 also slows the movement of keratinocytes in a wound-healing assay (30). Other studies have shown that BRG1 expression is downregulated by the suppressor of metastasis to the lung hepatic leukemia factor (HLF), and the genetic depletion of HLF or pharmacological degradation of BRG1 both suppress the migration of multiple cancer cell types in a collagen-coated environment (31). In these studies, the effect of BRG1 deletion on cell motility was attributed to alterations of the transcriptional events that BRG1 controls. Invasion in 3D matrices, apart from the mentioned nuclear deformability, also depends on actomyosin forces and proteolytic activity, particularly matrix metalloproteinases (32,33). As BRG1 and cohesin regulate gene expression, their inhibition may also influence cytoskeletal dynamics or MMP activity, in addition to altering nuclear stiffness. The present results suggest that in addition to these protein expression effects, physical changes in the nucleus immediately resulting from inactivation of BRG1 ATPase activity, even without removal of the protein or destruction of the BAF complex, can also alter cell motility.

Earlier studies showed no detectable effects on karyoplast stiffness by acute pharmacologic inhibition of DNA polymerase, RNA polymerase, or topoisomerase (4). Our current studies also show negligible acute effects of RNA polymerase or topoisomerase inhibitors on the migration of four different cancer cell types through pores (Figure 7), although longer-term inhibition of topoisomerase by doxorubicin leads to cell death (Figure 9). Inhibition of the nuclear ATPase cohesin, however, has significant effects on karyoplast stiffness and motility in several of the cell types studied, and the effect is the opposite of that of inhibiting BRG1; namely, cohesin inhibition makes the nucleus softer and enhances cell motility through small pores.

Although cohesin is a chromatin-binding ATPase, whether it functions as a traditional motor, like BRG1, with a defined step size and stall force, or as a molecular ratchet that rectifies motions produced by other forces, remains an area of active discussion. The SWI/SNF or BAF complex, which is powered by BRG1 or BRM, has been characterized by single-molecule methods to have a step size of 2 base pairs (∼7 nm) and a stall force of 30 pN. This corresponds to 5 kT of work, which can be accounted for by the 12 kT of free energy of a single ATP hydrolysis. In contrast, a recent study reports that cohesin has a step size of 10□nm and a stall force of 15□pN. That corresponds to 36 kT of mechanical work, which would require at least 3 ATP hydrolysis steps. The two ends of the structurally bivalent cohesin complex bind to two different DNA strands, either from different chromosomes, such as in the binding of sister chromatids, or from the same DNA strand to form chromatin loops within an interphase cell. The motor-like action of cohesin to extrude loops might require either thermal motions or fluid flow *in vitro* (34,35) or active movement caused by other motors within the nucleus (36), but in either case, cohesin would act like a crosslinker of DNA strands (37) in which ATP hydrolysis leads to the release of one of the DNA-binding sites (38). The peptide inhibitor of cohesin used in these studies inactivates its crosslinking effect, leaving cohesin tethered to chromatin but unable to link 2 strands together (39). This loss of crosslinking activity is consistent with the decrease in elastic modulus and increase in dissipation observed by global AFM deformation, and, at least in some cell types, also leads to an enhanced ability of cells to migrate through tight spaces in the transwell plate assay or into a collagen gel. Unlike BRG1 inhibition, which has similar effects on all the cell types tested here, the effects of cohesin inhibition on single-cell motility are cell-type dependent. There is a strong effect of increasing motility through the smallest pores for Huh7 and A549 cells, and a smaller enhancement for MCF7 cells, but the opposite effect on L18 cells. Whether the different effect in LN18 cells stems from possible mutations in cohesin or some other factor is not known. However, in the two cancer cell types that robustly made spheroids, cohesin inhibition permitted or enhanced outward migration from spheroids into the surrounding matrix for both LN18 and A549 cells.

In summary, the results presented here show that the activity of ATPases within the nucleus can have large effects on the global mechanics of the nucleus, with inhibition of specific ATPases either making the nucleus stiffer and more elastic or softer and more fluid. These effects on nuclear mechanics lead to equally large effects on the ability of cells to move through tight spaces that require significant deformation of the nucleus. The continuous active remodeling of chromatin within an interphase nucleus, therefore, appears to be required not only to organize the chromatin within the limited space of the nucleus or to transiently reveal DNA sequences for transcription but also to control the ability of the nucleus to change its shape in response to external mechanical stresses. The nucleus, therefore, is not adequately modeled as a viscoelastic polymer network surrounded by a membrane-like nuclear lamina, but rather as a continuously actively reorganizing heterogeneous system with both local reorganization driven by ATP-dependent motors and global shape changes driven by the same motors. This actively driven motion controls not only the chemical machinery within the nucleus but also the mechanical properties of the cell.

## Acknowledgements

This work was supported by the US National Institutes of Health and the US National Science Foundation through grants NIH R35GM136259 and NSF CMMI-1548571

## Author contributions

Ł.S., F.J.B., and P.A.J. designed the research. Ł.S., and F.J.B. performed the research. Ł.S., F.J.B., T.T.D., and P.A.J. analyzed the data. Ł.S., T.T.D., and F.J.B, performed image analysis. Ł.S., F.J.B., and P.A.J. wrote the paper.

