## Supplemental materials for "Role of nuclear ATPases in nuclear mechanics and cell migration through confined spaces: opposite effects of BRG1 and cohesin"

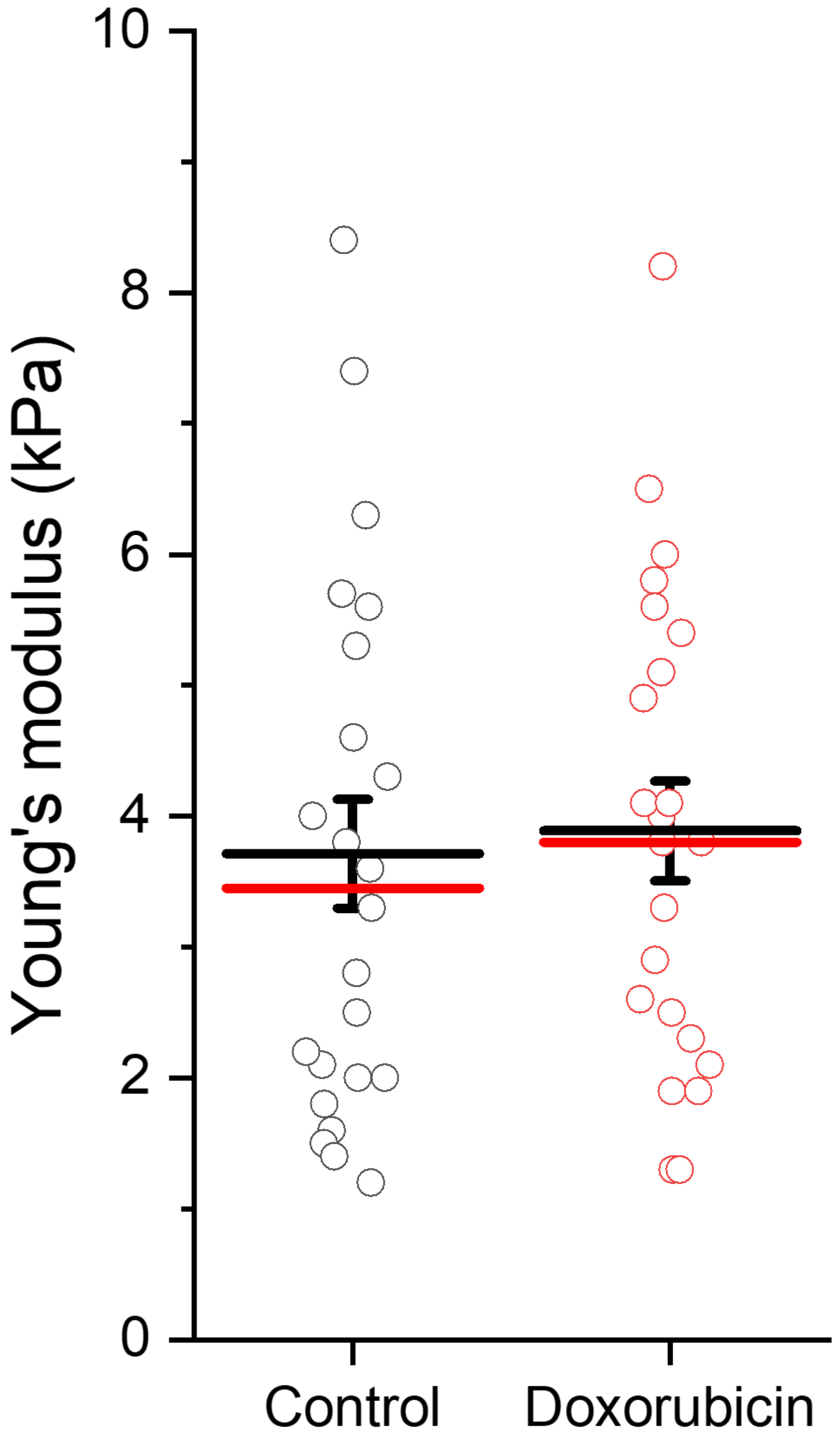


**Supplemental Figure 1.** Effect of doxorubicin (100 µg/mL) on the apparent Young's modulus of mEF -/- karyoplasts during a cycle of compression and recovery. Data are presented as mean ± SE from at least 3 independent experiments. No significant differences were detected (Student's t-test).

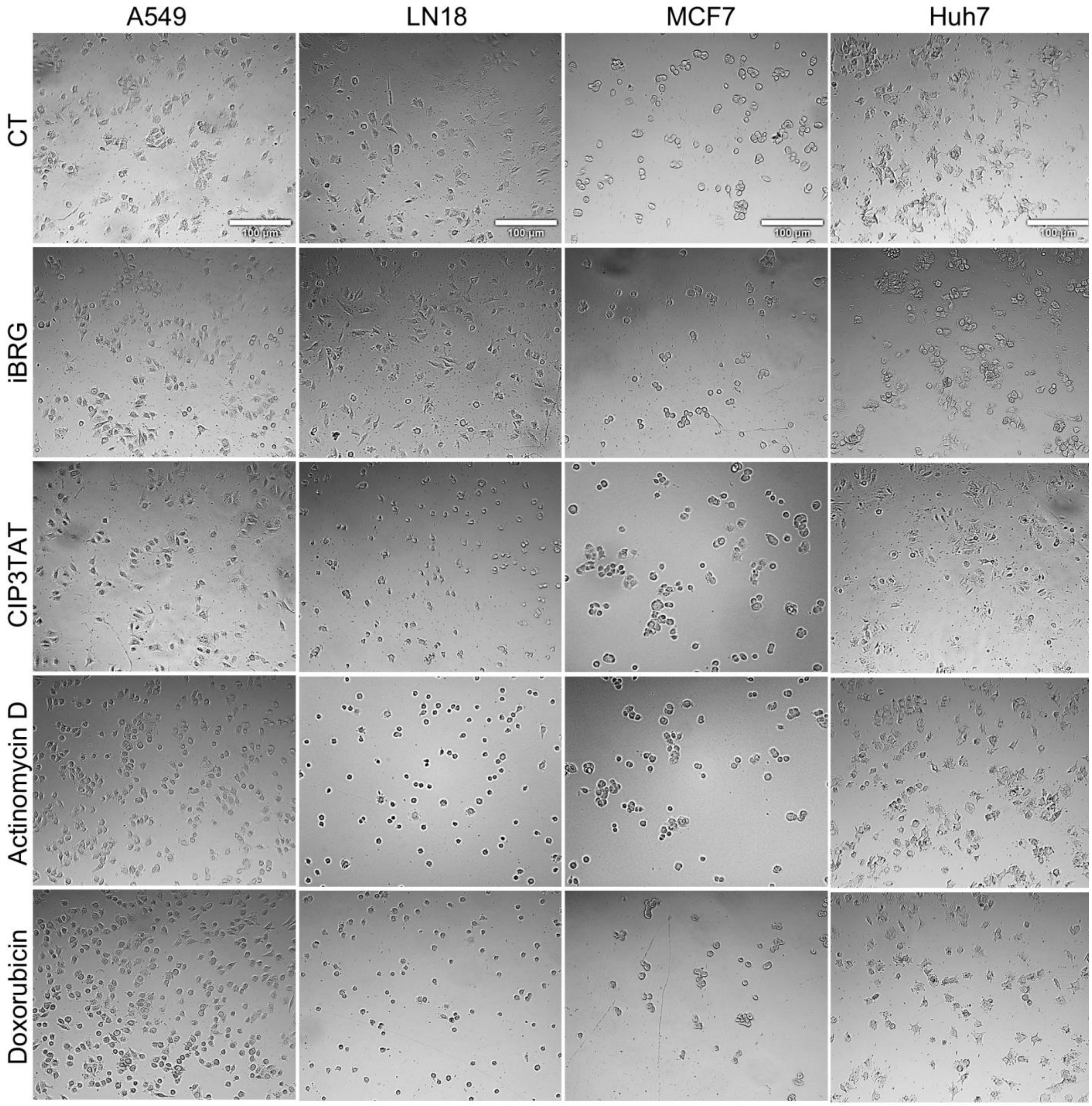


**Supplemental Figure 2.** Bright-field images of cancer cells (A549, LN18, MCF7, and Huh7) after 4 h treatment with DMSO (control, CT), iBRG (100 μM), CIP3TAT (10 μM), actinomycin D (100 μg/mL), or doxorubicin (100 μg/mL). Images are representative of cell morphology on the upper surface of Transwell membranes. Scale bar, ~100 μm.

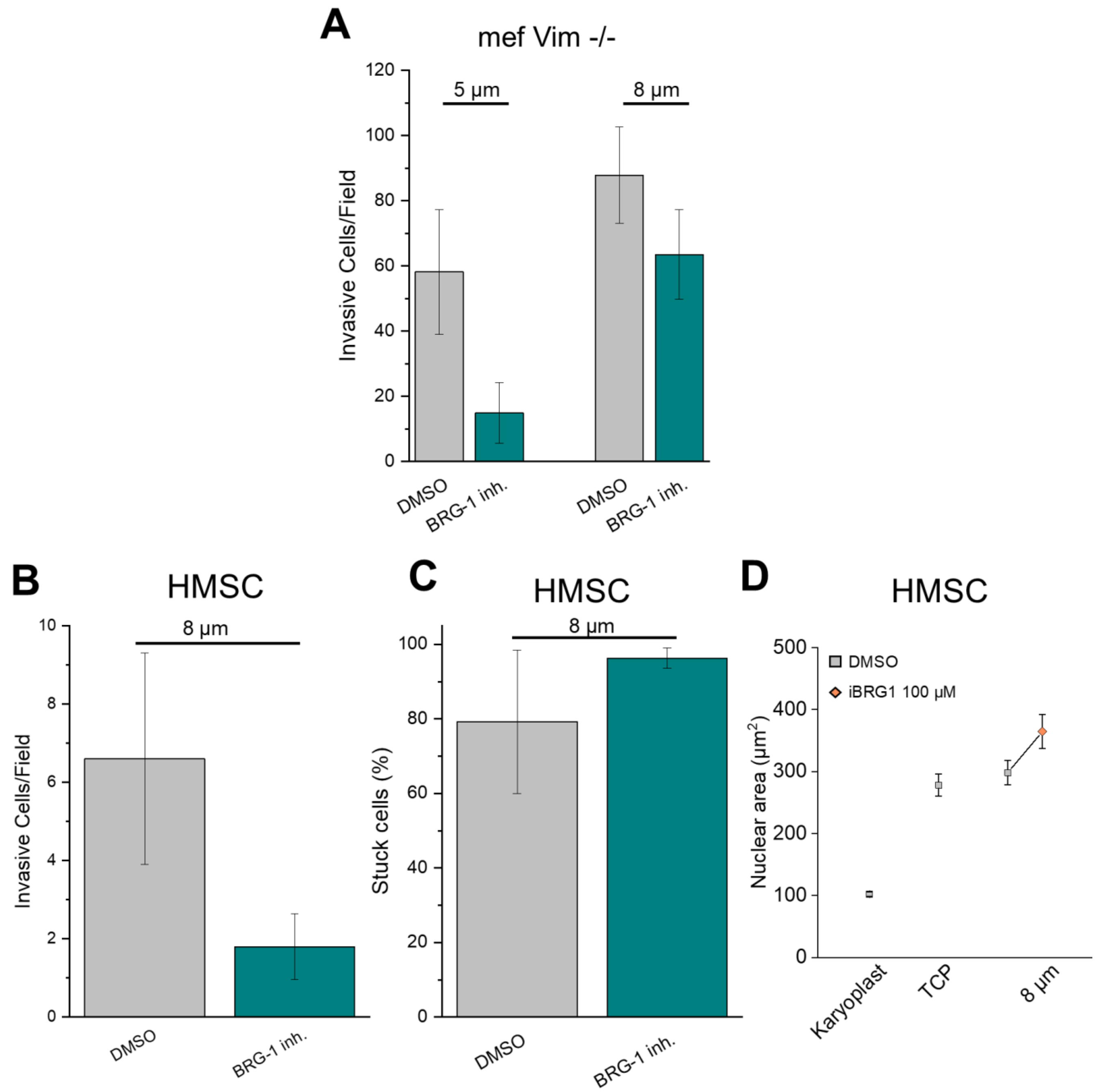


**Supplemental Figure 3.** Effect of inhibiting BRG1 (100 µM) on the migration of fibroblasts (**A**, mef -/-) and human mesenchymal stem cells (**B and C**, HMSC) through 5 and 8 micron-sized pores in the Transwell assay after 4 hours. (**D**) The projected nuclear area in cells that have passed through pores of various sizes, compared to the area of nuclei on flat surfaces before passage and within karyoplasts. Data are presented as mean ± SE; n ≥ 2 of independent experiments (5-10 images each).

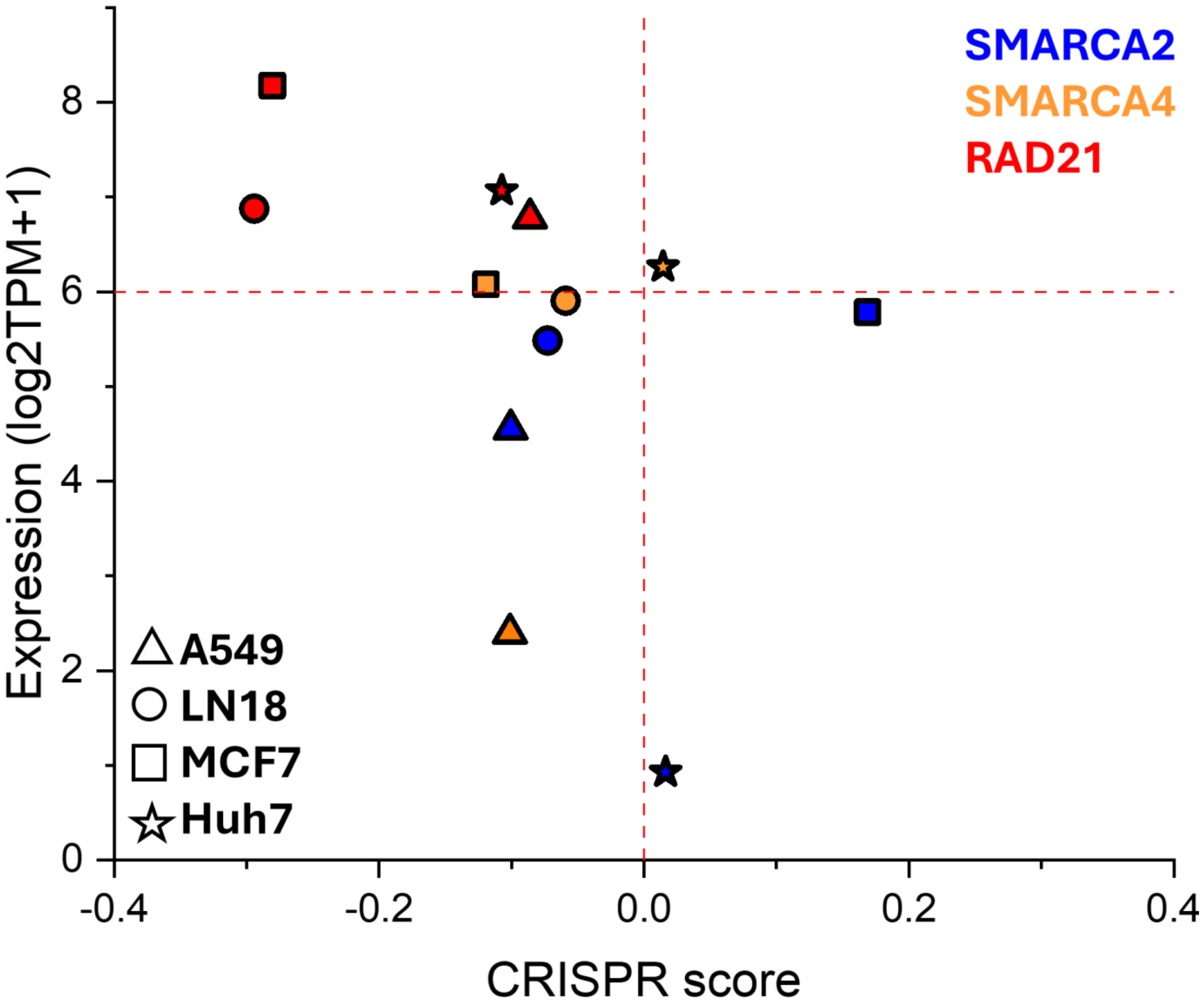


**Supplemental Figure 4.** RNA expression ($log_2$[TPM+1]) and CRISPR-Cas9 dependency (CRISPR scores) were obtained from the DepMap 25Q2 release. Values were extracted for A549, LN18, MCF7, and Huh7 cell lines, focusing on SWI/SNF (BRG1 – SMARCA4 and BRM – SMARCA2) and cohesin (RAD21). Expression values >6 were considered robust, while CRISPR scores <0 indicated essentiality.
